# Redundant function of Ets1 and Ets2 in regulating M-phase progression in post-natal angiogenesis

**DOI:** 10.1101/2020.02.19.956417

**Authors:** Sankha Ghosh, Catherine B. MarElia-Bennett, Blake E. Hildreth, Julia E. Lefler, Sudarshana M. Sharma, Michael C. Ostrowski

**Affiliations:** Department of Cancer Biology and Genetics and Comprehensive Cancer Center, The Ohio State University Wexner Medical Center, Columbus, Ohio 43210; Department of Biochemistry and Molecular Biology and Hollings Cancer Center, Medical University of South Carolina, Charleston, South Carolina 29425

**Keywords:** Ets1, Ets2, Genomics, G2/M transition, tumor-angiogenesis

## Abstract

Angiogenesis is a highly orchestrated process involving complex crosstalk between several endothelial cell processes including cell cycle, cell survival and migration. Transcription factors ETS1 and ETS2 are required for EC functions necessary for embryonic angiogenesis. Here, by utilizing endothelial cell-specific deletion of *Ets1* and *Ets2* on post-natal angiogenesis we demonstrate that ETS1 and ETS2 have a redundant regulatory role in controlling the expression of key G2/M regulators and anti-apoptotic gene modules by recruiting the transcriptional activator CBP/p300 to the enhancers of these genes. Further, cultured aortic ECs lacking both ETS1 and ETS2 demonstrated G_2_/M phase arrest and increased apoptosis. *In vivo*, EC infiltration and invasion was attenuated when matrigel admixed with mouse mammary tumor cells was injected into adult mice with EC-specific ablation of *Ets1* and *Ets2*. These results demonstrate that deletion of *Ets1* and *Ets2* in endothelial cells inhibits angiogenesis by altering cell cycle progression and decreasing cell survival.

---

ETS1 and ETS2 belong to a family of transcription factors that share a highly conserved helix-winged-helix DNA-binding domain and a core DNA-binding consensus sequence 5’-GGA(A/T)-3’ motif (1). ETS family factors influence a wide range of cellular processes and can act as transcriptional repressors and activators depending on the target gene and post-translational modifications (1). For example, ETS factors, including ETS1 and ETS2, have an integral role in the development and function of endothelial cells (ECs). Nearly all EC enhancers and promoters characterized to date contain an ETS binding site, many of which can be bound by multiple ETS factors (2). The blood vascular network serves as the principal communication channel between various organs and tissues of the body. It is also central to maintaining homeostasis and coordinating wound repair. Thus angiogenesis, the formation of new blood vessels from a preexisting vascular network, is essential for embryonic development and normal physiology as well as pathological conditions including tumorigenesis (3–6). Once a blood vessel network is formed in the adult, the differentiated endothelial cells (ECs) lining the inner surface of mature blood vessels adopt a quiescent, G_0_-like state. However, these cells remain highly plastic and in response to angiogenic factors, ECs re-enter cell cycle and proliferate, a pivotal step in neo-angiogenesis (7).

Our previous studies demonstrated that EC specific deletion of *Ets2* in combination with global *Ets1* knockout results in defective angiogenesis and embryonic lethality. This phenotype can be rescued by the presence of a single copy of either *Ets1* or *Ets2*, indicating the redundant nature of their functions (8). Migration and apoptosis were EC processes affected by the absence of *Ets1* and *Ets2* through the dysregulated expression of genes like *Bcl2l1* and *Mmp9*. However, the lethal embryonic phenotype at e11.5-e12.5 prevented a detailed analysis of mechanisms underlying the observed vascular defects (8).

We conducted the current study to further define the mechanism by which the overlapping functions of ETS1 and ETS2 affect postnatal EC biology and function. Combining global gene expression and chromatin immunoprecipitation sequencing (ChIP-seq) using primary aortic EC cultures, we identified novel gene targets, including *Ube2c*, *Cdc25c* and *Birc6*, and a previously unknown function for ETS1 and ETS2 for regulating M-phase of the cell cycle. Further, genetic analysis of *Ets1* and *Ets2* in tumor angiogenesis provided evidence for an EC-autonomous, redundant function of these factors.

## Results

### Loss of ETS1 and ETS2 in ECs result in deregulation of genes critical for cell survival and proliferation

Previous work from our group demonstrated that mice with genotype *Tie2-Cre*;*Ets1*^−/-^;*Ets2*^*fl/fl*^ are not viable beyond E14.5 due to aberrant angiogenesis (8). To better address the underlying mechanism responsible for the angiogenic defects observed in the double mutant mice, we generated a conditional *Ets1* allele by flanking exons 7 and 8 with *loxP* sites (Figure S1A), as deletion of these exons encoding the DNA binding domain produce a null allele (9). A tamoxifen-inducible transgene *Tie2-Cre-ER*^*Tam*^ (10) was used to affect conditional deletion of the *Ets1*^*loxP*^ allele and the previously characterized *Ets2*^*loxP*^ conditional allele (8). Using this inducible *Tie2-Cre* we could demonstrate recombination of both *Ets1*^*loxP/loxP*^ and *Ets2*^*loxP/loxP*^ alleles in aortic ECs isolated from 6-week-old mice homozygous for both *Ets* alleles following tamoxifen treatment, resulting in decreased mRNA expression of both genes (Figure S1B, C).

Expression profiling with the Affymetrix platform was performed using RNA isolated from low-passage, primary murine aortic ECs from each of four genetic groups: wild-type, *Ets1* KO, *Ets2* KO and *Ets1/Ets2* DKO. The EC populations were purified from the aorta of mice of appropriate genotype, and Cre was delivered by lentivirus to affect conditional gene deletion (Figure S1C). The expression data obtained was analyzed with Gene Set Enrichment Analysis (GSEA) comparing the expression pattern of the DKO cells with that of the other three genotypes combined (11). The analysis identified ~2,000 genes differentially expressed in the DKO cells versus the other genotypes; the top 200 genes differentially expressed in DKO compared to the three controls are represented in the heat map in Figure 1A. Using stringent criteria (nominal p-value < 0.01, FDR q value < 5%), GSEA identified only three fundamental cellular processes affected in ECs by the deletion of *Ets1* and *Ets2*: M-phase of the cell cycle, cell survival, and cell migration (Figure 1B and Figure S1D). Genes at the leading edge of the gene sets identified by GSEA were prioritized for further analysis. Using qRT-PCR with RNA prepared from independent aortic EC samples, the expression of three genes representing the mitotic M-phase gene set (*Cdc25c*, *Anapc5*, and *Ube2c*) and three representative anti-apoptosis genes (*Birc6*, *Bcl2l1*, and *Sphk1*) were analyzed in cells with the four genotypes. The results confirmed that loss of either *Ets1* or *Ets2* alone did not affect the expression of these genes. In contrast, in DKO ECs lacking both *Ets1* and *Ets2*, the expression of both the M-phase and anti-apoptosis genes was downregulated 3-to 15-fold compared to the wild-type counterparts (Figure 1C, D).

**Figure 1.**
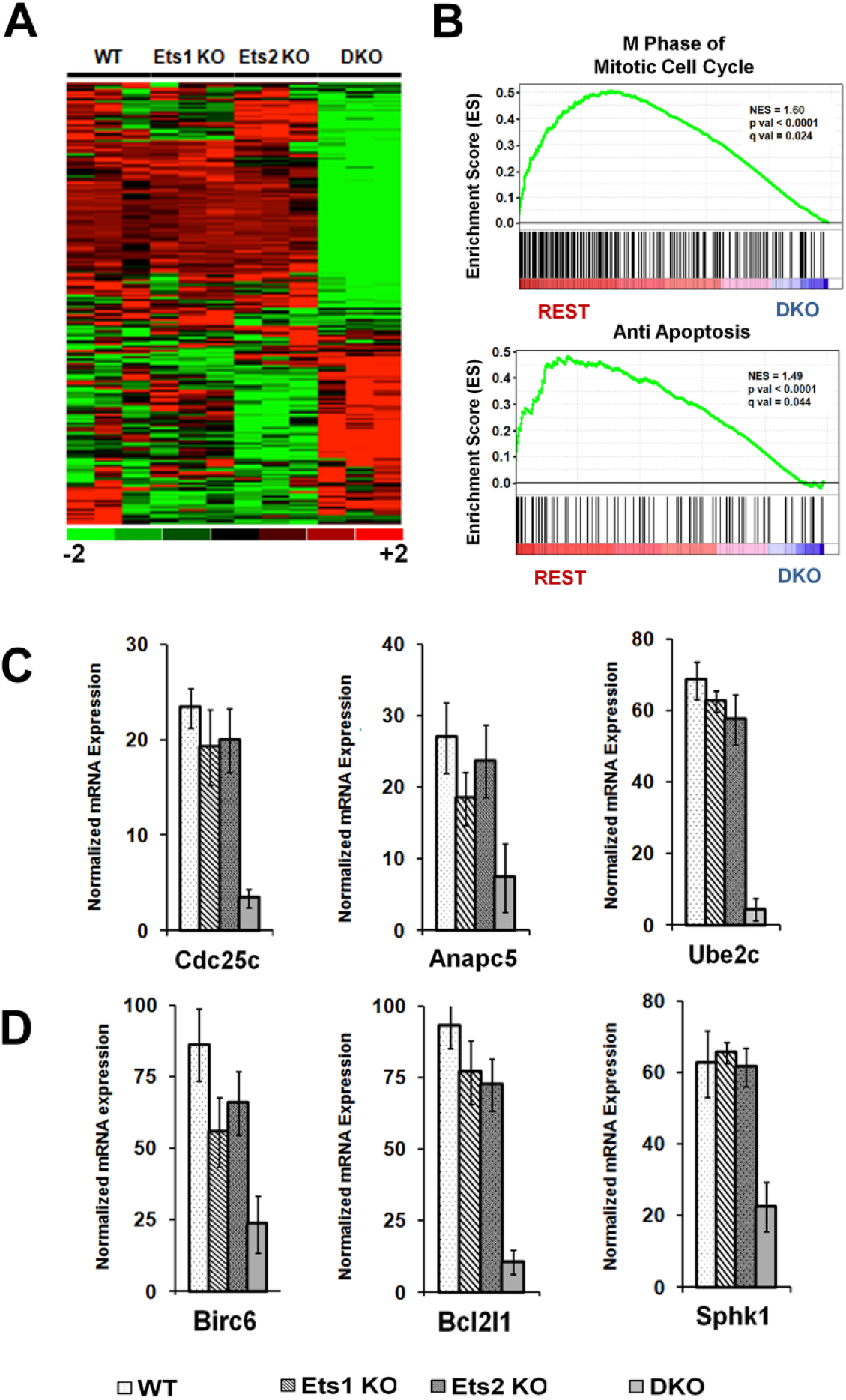
Deletion of Ets1 and Ets2 Results in Deregulation of Genes Involved in Cell Cycle and Cell Survival. A). Heat map of top 200 genes dysregulated in Ets1/Ets2 double knockout endothelial cells compared to WT, Ets1 single knockout, Ets2 single knockout endothelial cells. B). GSEA analysis representing top two enrichment categories enriched in absence of both Ets1 and ETS2 compared to WT, ETS1KO, ETS2 KO endothelial cells. C). Independent qRT-PCR analysis of genes critical for M-Phase from WT, Ets1KO, Ets2KO and ETS1/2 DKO endothelial cells. D). Independent gene expression analysis of anti-apoptotic genes in the endothelial cells lacking either ETS1, ETS2, or both ETS1 and ETS2 compared to WT endothelial cells. Data in C and D are the average of three independent experiments done in duplicate. Error bars represent the standard deviation. Only ETS1/2 DKO endothelial cells were statistically different from the other three genotypes (p<0.05).

### Enhancers occupied by both ETS1 and ETS2 specifically regulate G_2_/M-phase transition and cell survival genes

In a complementary analysis, the genome-wide distribution of ETS1 and ETS2 binding sites were determined in wild-type aortic ECs by chromatin immune-precipitation coupled with massively parallel sequencing (ChIP-seq). The resulting sequence reads were aligned to the reference mouse genome (mm10) (12) with Bowtie2 (13), and subsequently analyzed with the HOMER suite of software (14). The analysis identified 77,287 ETS1 binding peaks and 63,154 ETS2 binding peaks, with 12,919 of these genomic regions bound by both transcription factors (Figure 2A, B and Supplementary Table 1). Of the 77,287 ETS1-bound peaks, 7% of the loci were associated with promoter-TSS regions with the remaining peaks at distal enhancers (Figure 2C, S2A). Genomic distribution of ETS2-bound regions mirrored the ETS1 distribution with a majority of distal enhancers and 9% of promoter-TSS loci (Figure 2B, Figure S2E). In contrast, loci with ETS1/ETS2 co-bound peaks were comparatively more abundant in promoter-TSS regions (16%; Figure 2E).

**Figure 2.**
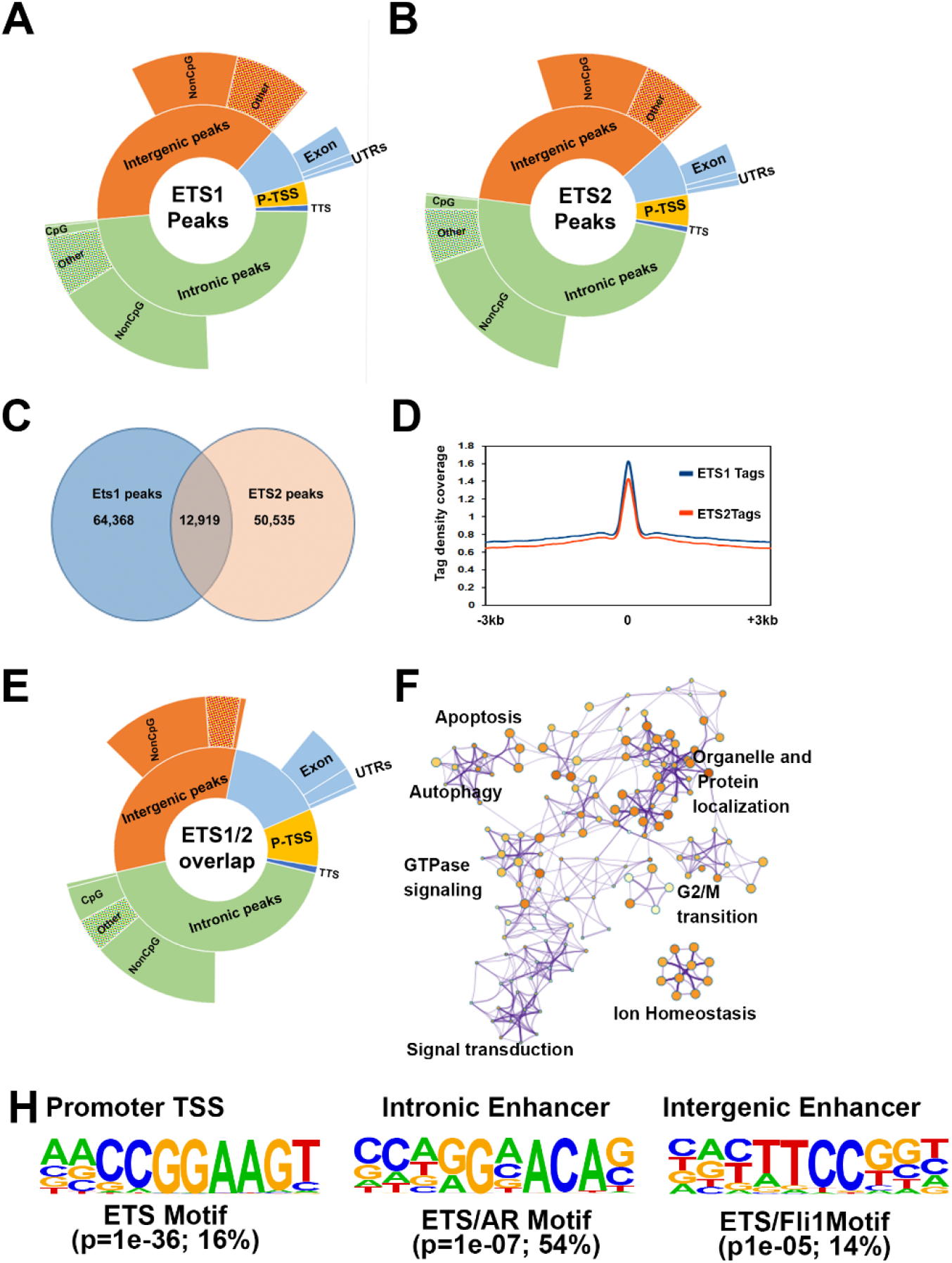
ETS1 and ETS2 bound distal enhancers are associated with genes responsible for G2/M transition and cell survival. A) Graphical representation of the distribution of ETS1 ChIP-Seq peaks throughout the genome in endothelial cells. B) Graphical representation of the distribution of ETS2 ChIP-Seq peaks throughout the genome in endothelial cells. C) Graphical representation of the distribution of ETS2 ChIP-Seq peaks throughout the genome in endothelial cells. D) ETS1 and ETS2 ChIP-Seq tag density coverage around ETS1/ETS2 co-bound peaks. E) Graphical representation of the distribution of ETS1/ETS2 co bound peaks in the genome. F) Pathway enrichment analysis of ETS1/ETS2 co bound gene loci demonstrating enrichment of genes responsible for G_2_/M transition and Apoptosis. G) Ranked enriched Motifs from ETS1 and ETS2 co-bound promoters, intronic enhancers and Intergenic enhancers, both percentage of sequences containing the motif and statistical significance were considered.

The regulatory elements enriched for both ETS1 and ETS2 were associated with genes by nearest neighbor analysis and overlapped with genes differentially expressed in *Ets1/Ets2* DKO ECs compared to wild-type ECs. Motif analysis indicated that the promoter regions in ETS1-bound, ETS2-bound or ETS1/ETS2 co-bound regions predominantly exhibit ETS consensus binding motifs in approximately 20% of the loci. However, the distal enhancers tend to exhibit an ETS-AR half site in the majority of the peaks analyzed (Figure 2H, Figure S2D, H).

When genomic loci occupied by both ETS1 and ETS2 were analyzed for the enrichment of consensus motifs (see Materials and Methods), as expected the occupied promoter-TSS loci contained predominantly ETS binding sites, but other transcription factor motifs were not significantly enriched (Figure 2H). However, motif enrichment analysis of enhancers bound by either ETS1 alone or ETS2 alone provided insight into potential cooperative partners such as androgen receptor (AR) and Tal1, which are both involved in angiogenesis (Figure S2D, H) (15,16). The loci enriched for ETS1 or ETS2 alone were independently overlapped with WT/*Ets1* KO and WT/*Ets2* KO gene expression data, respectively. This analysis indicated that apoptosis was the only common gene ontology (GO) term shared between DKO cells and single KO cells (Figure 2F and Figure S2C, G, respectively). Single KO cells shared a few pathways, (e.g., growth factor/TGFβ signaling) but the majority of the GO terms were distinct (Figure S2C, G).

We next focused on the differentially expressed genes with linked ETS1/ETS2 binding sites that were represented in the biological processes identified by the GSEA analysis (Figure 1). A significant fraction of differentially expressed M-phase genes (55%) and cell survival genes (58%) were co-bound by ETS1 and ETS2. The same six representative genes involved in regulating M-phase (*Cdc25c*, *Anapc5*, and *Ube2c*) or in promoting cell survival (*Birc6*, *Bcl2l1*, and *Sphk1*) were studied in Chromatin immunoprecipitation (ChIP) assays using ECs of the same four genotypes as above. Both ETS1 and ETS2 were found to be enriched at the regulatory regions of the genes examined in the wild-type chromatin. In single KO cells, the absence of either transcription factor alone did not significantly reduce the binding of the other (Figure 3). However, the association was completely abolished when neither ETS1 nor ETS2 was present (Figure 3; fourth bar graph in each panel).

**Figure 3.**
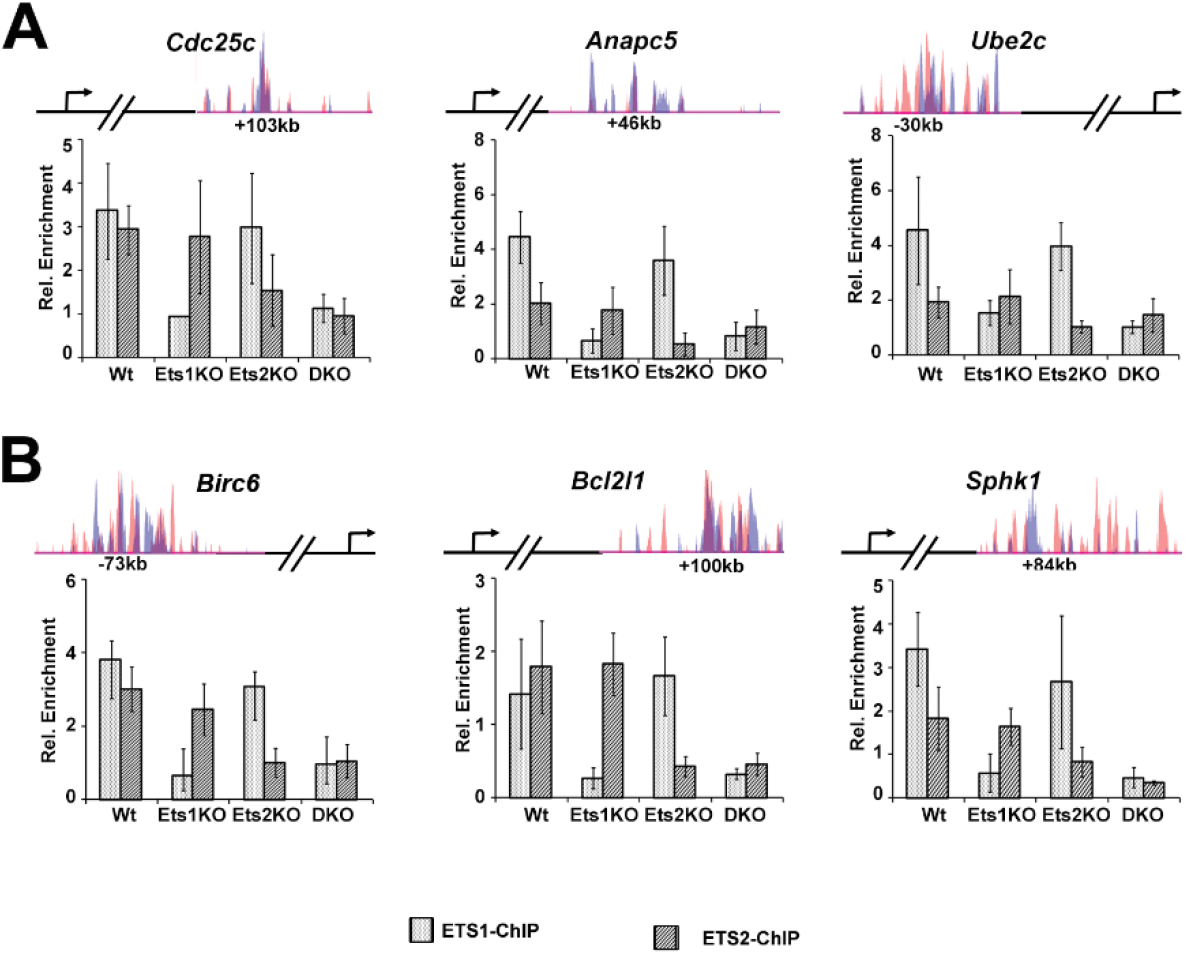
Conventional ChIP followed by qPCR confirms binding of ETS1 and ETS2 distal enhancers are associated with genes responsible for G2/M transition and cell survival. A and B) Top panel: Overlay of Ets1 and ETS2 co-bound ChIP-Seq peaks that were verified by conventional ChIP followed by qPCR (bottom pnel) for genes important for M phase of cell cycle (A) and cell survival (B) (bar graphs, n=3) Data represented as Average of three independent experiments, error bars represent SD. Non-normally distributed data and/or data with unequal variance underwent logarithmic transformation. All data were then available for comparison with a one-way ANOVA and Holm-Sidak post-hoc analysis between select groups.

### Presence of either ETS1 or ETS2 is sufficient for the recruitment of transcriptional coactivator CBP/p300 complex to loci governing the expression of genes necessary for endothelial cell proliferation and survival

The CREB-binding protein (CBP) and p300 are two highly homologous nuclear proteins that function as transcriptional coactivators by acting as adapters between transcription factors and basal transcription machinery (17). Mitogen-activated protein (MAP) kinase induced phosphorylation of ETS1 and ETS2 leads to recruitment CBP/p300 to enhance transcriptional activation (18,19). In order to determine whether CBP/p300 mediates the regulation of the selected genes co-bound by ETS1 and ETS2 globally, CBP-p300 ChIP-seq data from murine models (20) were overlapped with ETS1/ETS2 co-occupied peaks. Relative tag densities of combined CBP/p300 ChIP-seq data from endothelial cells from whole embryo, embryonic heart, and embryonic lung (20) were over layered with ETS1/ETS2 co-occupied peaks using K means clustering (21,22) followed by Treeview visualization algorithm (23,24). In concurrence with previous work on ETS1 and CBP/p300 association in Jurkat T-cells (25), we observed CBP/p300 tags were highest around the ETS1/ETS2 co-occupied peaks in the promoter-TSS regions, followed by distal intergenic or intronic enhancers (Figure 4A). We analyzed CBP/p300 recruitment at the loci of the same six representative genes in ECs derived from each of the four genetic groups. While CBP/p300 recruitment was observed when either ETS1 or ETS2 was present, with the exception of *Sphk1*, the absence of both factors abolished CBP/p300 recruitment (Figure 4B, C).

**Figure 4.**
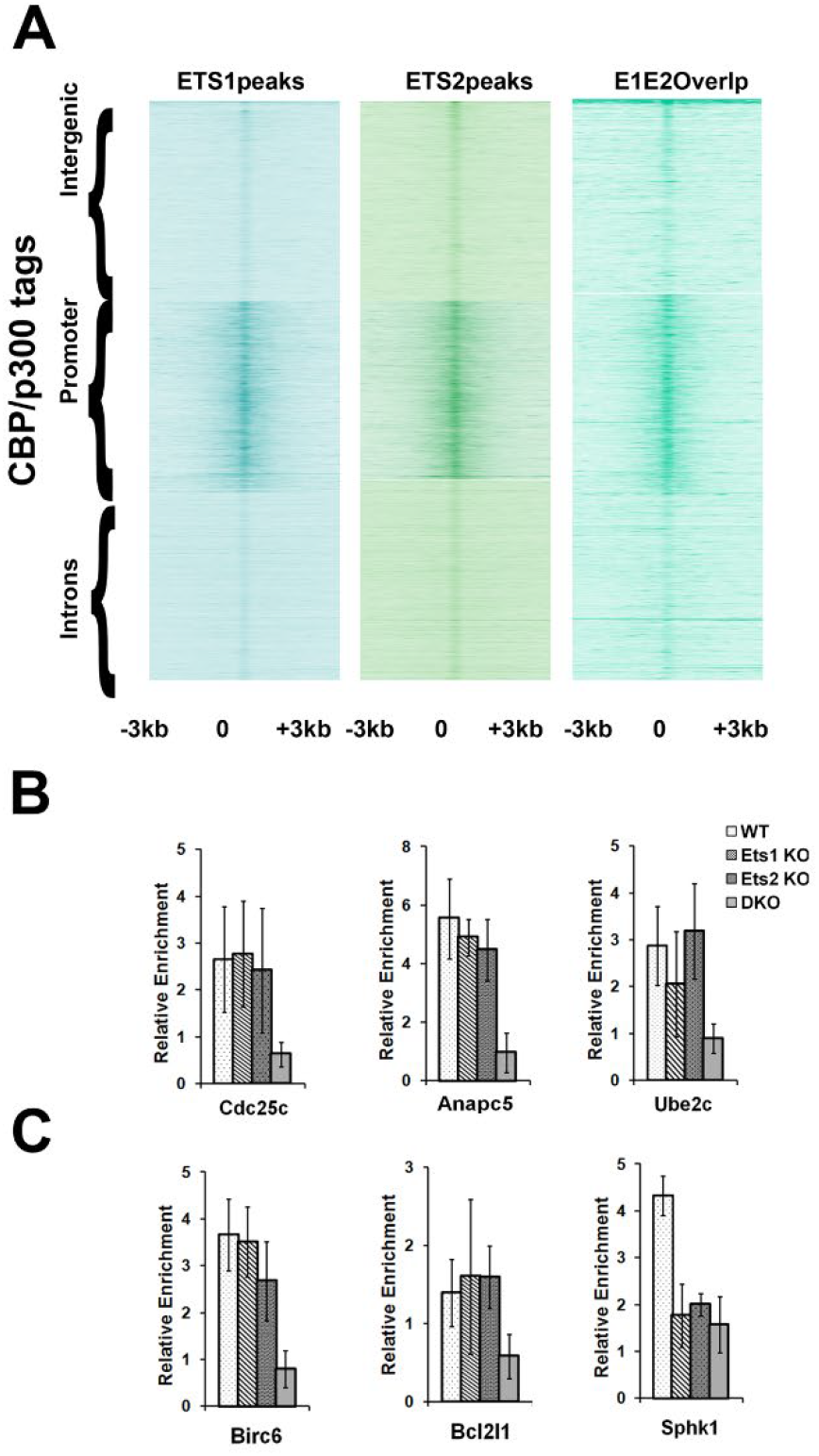
ETS1 and or ETS2 is required for the recruitment of acetylated CBP to the enhancer regions of genes governing G2/M transition and cell survival. A) Treeview plot of tag densities of combined CBP/p300 ChIP-seq data flanking ± 3kb region of either ETS1(left panel), or ETS2 (middle panel), and co-bound ETS1-ETS2 ChIP-seq peak centers (right panel). B) Conventional ChIP followed by qPCR analysis of ETS1 and ETS2 co-bound peaks for the association of acetylated CBP/p300 occupancy on Mitotic genes represented in Figure 3. C) ChIP-qPCR verification of anti-apoptotic genes using acetylated CBP/p300 antibody as described in Figure 3.

### Deletion of ETS1 and ETS2 results in G_2_/M-phase cell cycle arrest and induction of apoptosis in endothelial cells

To further test the hypothesis that ETS1 and ETS2 regulate cell cycle and apoptosis, ECs derived from the four ETS genotypes were subjected to propidium iodide (PI) staining followed by flow cytometry. A normal distribution of the cell population throughout the cell cycle was observed in each of the control EC groups (Figure 5A). Of note, there was a ~3-fold increase in the G_2_/M-phase population in the DKO cells compared to the controls (Figure 5A), indicating cell cycle arrest at the G_2_/M transition. To validate this result, ECs were subjected to phospho-histone H3 staining. Histone H3 is phosphorylated during mitosis and can be used as a specific immunomarker for cells undergoing mitosis (26). The results of this experiment showed a significant increase in the mitotic index (MI) in the DKO cells. In control cells, the MI ranged between 8 to 10% while the MI of the DKO cells was ~35% (Figure 5B).

**Figure 5.**
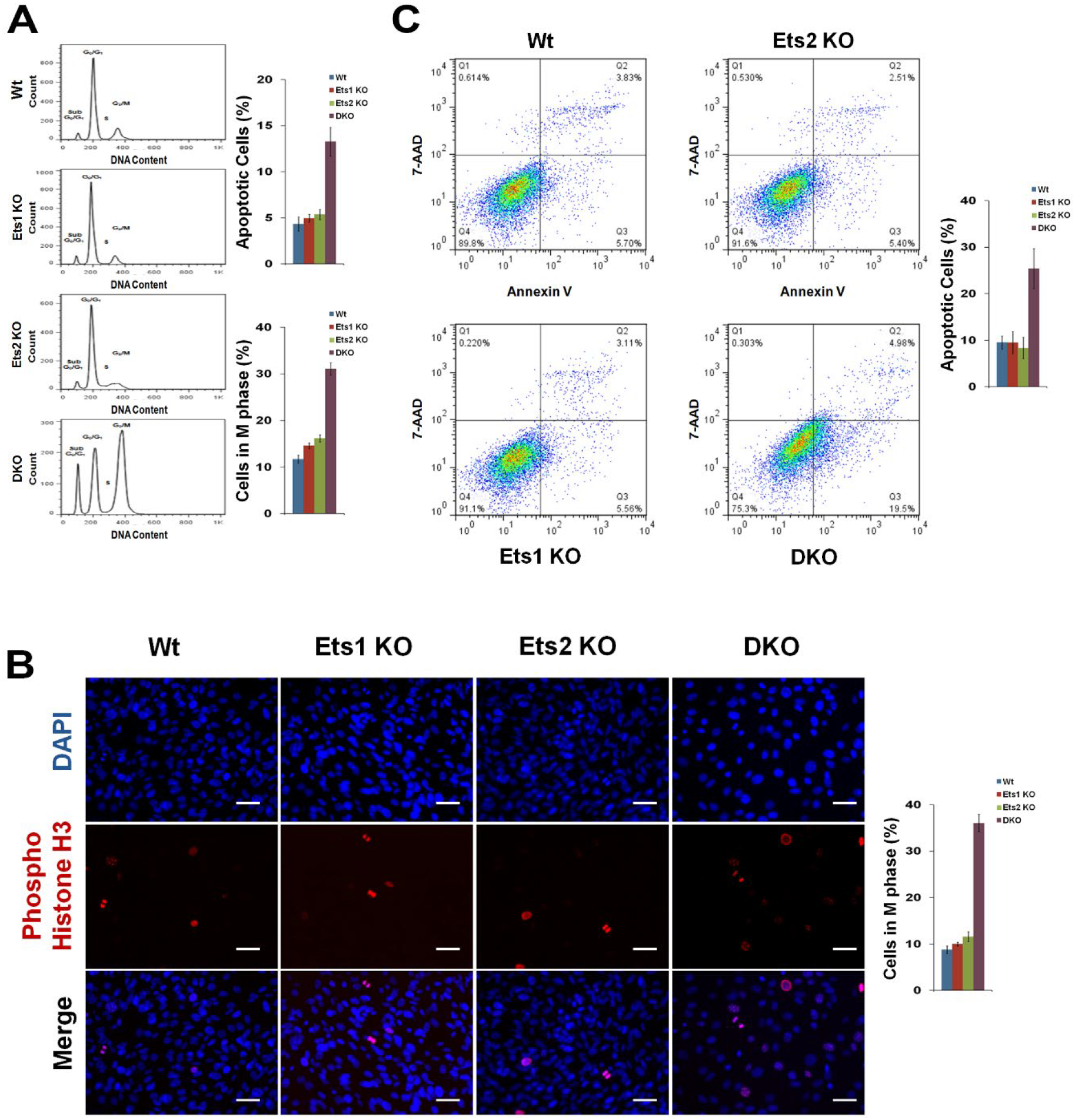
Ets1 and Ets2 are required for endothelial cell survival and cell cycle *in vitro.* A) Cultured aortic ECs were serum starved (0.1% FBS) for 24 h. Cells were then harvested and subjected to PI staining followed by FACS. Graphs at right summarize the quantification. The percent apoptosis and M phase is calculated from the subsequent sub-G_0_/G_1_ peak and G_2_/M peak as determined by cell-cycle analysis. Data are plotted as mean percentages ± SD of three independent experiments. B) Assessment of apoptosis after serum starvation (0.1% FBS, 24 h) of cultured aortic ECs by Annexin V/7-AAD staining followed by FACS analysis. ECs in the lower right quadrant are Annexin-positive, early apoptotic cells. The cells in the upper right quadrant are Annexin-positive/7-AAD-positive, late apoptotic cells. Graphs at right demonstrate the quantification. Data are plotted as mean percentages ± SD of Annexin V positive cells (both early and late apoptosis) of two independent experiments. C) Cultured aortic ECs were processed for indirect immunofluorescence using anti-phospho-histone-H3 antibody (red), DAPI was used to stain the DNA. The graphic panel at right shows the number of M phase cells for each genotype expressed as mean percentages ± SD of the total number of cells measured from two independent experiments. Scale bars: 50 um (C)

In addition to an increase in G_2_/M cells, the fraction of DKO cells in the sub-G_0_/G_1_ population was ~3-fold higher than in the controls. Moreover, ECs with only ETS1 or ETS2 demonstrated no significant changes in the number of cells in the sub-G_0_/G_1_ population compared to wild-type cells (Figure 5A). The results of the PI staining assay were confirmed by annexin V and 7-aminoactinomycin D (7-AAD) staining. Similar to the results of the PI staining, a ~2.5-fold increase in the number of cells in the apoptotic population was observed in the absence of both ETS1 and ETS2 (Figure 5C).

### Defective tumor angiogenesis upon deletion of ETS1 and ETS2 in endothelial cells

To better define the mechanisms underlying the function of the two ETS factors in tumor angiogenesis, we studied the effect of conditional deletion of *Ets1* and *Ets2* in adult mice using Matrigel plug assays. For this purpose, 10-week-old inducible EC-specific *Ets1*/*Ets2* double knockout (*Tie2-Cre-ER*^*Tam*^;*Ets1*^*fl/fl*^ *Ets2*^*fl/fl*^) mice and controls containing one copy of *Ets1* (*Tie2-Cre-ER*^*Tam*^;*Ets1*^*+/fl*^ *Ets2*^*fl/fl*^) were treated with tamoxifen and injected subcutaneously with Matrigel plugs containing MVT1 mouse mammary tumor cells (27). Robust growth of new CD31-positive blood vessels into the Matrigel plugs occurred in the control group that contained a single wild-type copy of ETS1 (Figure 6A). Notably, the number of vessels and degree of vessel branching were significantly decreased in the DKO mice (Figure 6A). In parallel studies, an *in vitro* Matrigel tube formation assay was performed with *Ets1/Ets2* DKO and control aortic ECs (Figure 6B). These assays demonstrated that the control ECs, including *Ets1* KO and *Ets2* KO cells, formed multicellular tubular structures 24 hours after plating while DKO cells failed to form the mature tube-like structures (Figure 6B).

**Figure 6.**
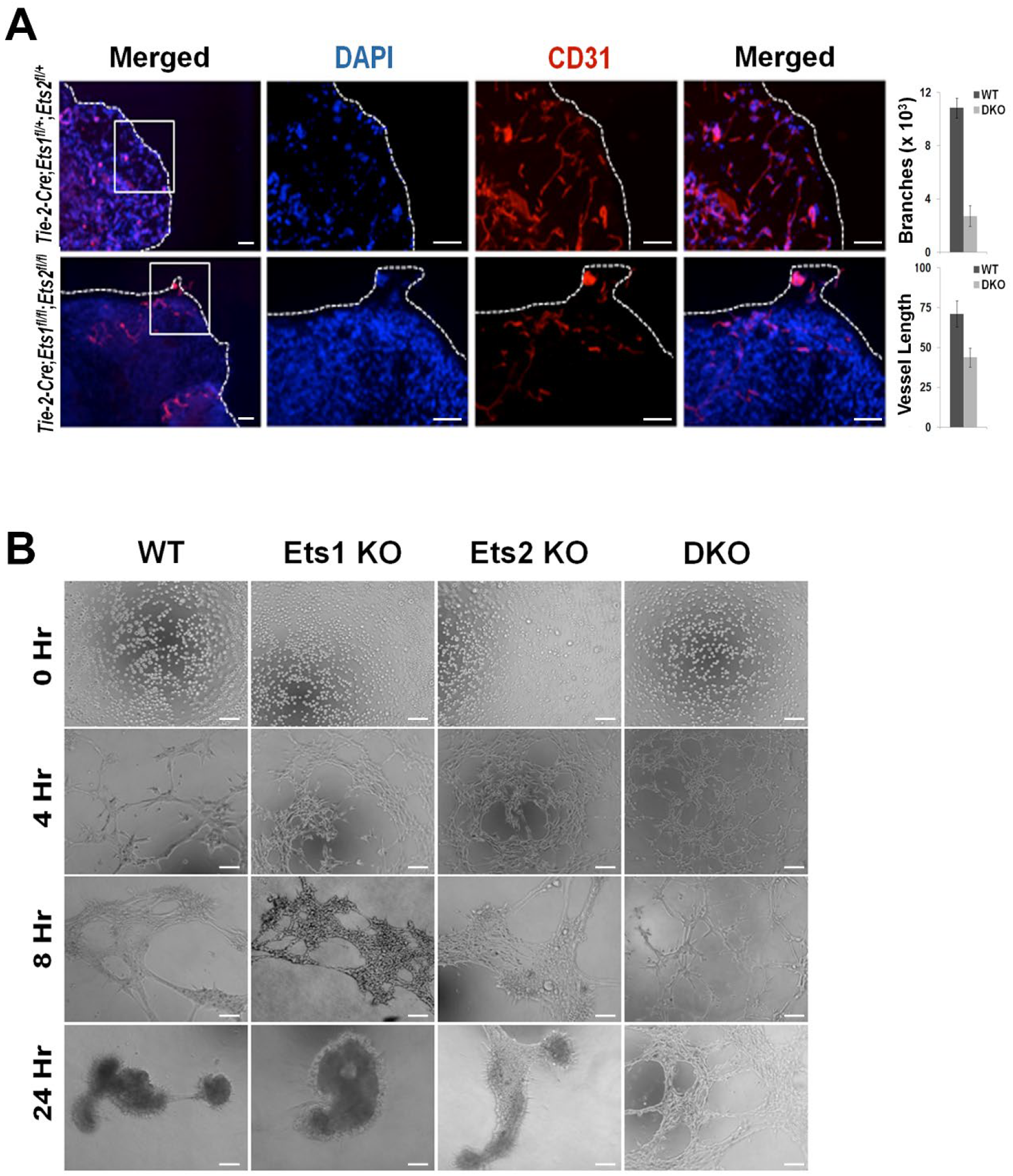
Ets1/2 are required for tumor angiogenesis. A) Staining for CD31 in matrigel plugs: Matrigel, with 5×10^5^ MVT1 cells, was subcutaneously injected into control and DKO mice and harvested after 10 days. Dashed lines indicate the outer margin of the plug. Region in the boxed area is magnified in the next panel of the corresponding genotype. Quantitation of CD31-positive vessels/field and branching is shown in the right panel. n=6 plugs B) *In vitro* matrigel tube formation assay: WT, Ets1 KO, Ets2 KO and DKO ECs were plated onto polymerized matrigel layer in a 96-well plate. Tube formation was monitored at 4 hours interval for 24 hours. Scale bars: 100 um left most panel, 50 um right three panels (A); 80 um (B).

## Discussion

Sprouting angiogenesis is a multi-step process that involves the orchestrated execution of discrete biological steps including basement membrane degradation, EC proliferation, directed EC migration, EC tube formation, and vessel fusion and pruning (3). The execution requires complex and highly coordinated crosstalk between angiogenic inducers and inhibitors, which engage specific transcription factors that modulate the biological processes (4,6,28). Defining the links between specific factors and EC processes is essential for revealing molecular details that regulate angiogenesis. In the current study, we demonstrate an unexpected function of ETS1 and ETS2 in controlling the M-phase of the endothelial cell cycle through regulating genes required for G_2_/M transition. Additionally, we extend the results of our previous studies demonstrating the role of ETS1 and ETS2 in preventing EC apoptosis by defining direct anti-apoptotic target genes. ETS1 and ETS2 act redundantly and in a cell autonomous manner to regulate cell cycle and apoptosis in ECs *in vitro* and tumor angiogenesis in a matrigel plug assay *in vivo*.

Data from previous investigations of the tumor microenvironment by our group have provided insights into how ETS2 may affect tumor growth and angiogenesis in a cell type-specific manner. For example, deletion of *Ets2* in the stromal fibroblasts of mouse mammary tumors resulted in reduced tumor growth and aberrant angiogenesis, whereas loss of *Ets2* in epithelial tumor cells had no significant effect (29). Additionally, an examination of the role of ETS2 in tumor associated macrophages (TAMs) revealed that ETS2 repressed the expression of anti-angiogenic inhibitors in mouse mammary tumors. Indeed, *Ets2* deletion in the TAMs resulted in reduced tumor growth, metastasis, and angiogenesis (27).

Interestingly, the redundant functions of ETS1 and ETS2 appear to be regulated from enhancers as opposed to promoter regions as the majority of the co-occupied peaks were found in the inter-and intragenic regions through recruitment of the CBP/p300 complex. In addition, standard ChIP performed on single KO cells confirmed that binding of one protein to these regulatory regions is completely independent of the other. Taken together, these data suggest the direct modulation of genes specific to cell cycle progression and cell survival by ETS1 and ETS2 is critical in their regulation of EC function. Notably, most of the ETS1/ETS2 co-bound regions were enriched for motifs of androgen receptor (AR) and E-box (Tal1) which are known effectors of angiogenesis.

Extracellular signal-regulated kinases 1 and 2 (ERK1 and ERK2) are upstream regulators of ETS1 and ETS2. Previous studies from our lab have shown that *Erk1/Erk2* ablation in ECs results in reduced EC proliferation. Additionally, global gene expression analysis comparing control and *Erk1/Erk2-*deleted ECs revealed dysregulation of a number of genes involved in cell cycle (30). Based on these findings, we hypothesized that the effects of Erk1 and Erk2 on endothelial cell cycle progression could be mediated by ETS1 and ETS2. Indeed, in our current study, ablation of ETS1 and ETS2 in ECs resulted in abnormal cell cycle and defective postnatal angiogenesis both *in vivo* and *in vitro*. The *in vitro* assays performed here demonstrate that the cell cycle was highly compromised in the *Ets1/Ets2* DKO cells. This led to an increased population of DKO ECs in the G_2_/M-phase, disrupting the normal cell cycle phases that are critical for angiogenesis. Furthermore, the deletion of *Ets1* and *Ets2* promoted endothelial cell death, indicating that apoptosis is the likely exit pathway for the mutant cells following arrest at M-phase.

Consistent with the above results, the GSEA analysis identified the M-phase of the cell cycle and apoptosis among the most important downstream effector processes of ETS1 and ETS2. Several genes involved in cell cycle regulation were differentially expressed in the DKO ECs which could account for the observed M-phase arrest. Similarly, the expression of a number of anti-apoptotic genes was also downregulated in the DKO cells, conferring increased sensitivity to apoptotic signals. Findings from the global DNA-protein interaction study suggested that several of these genes were direct targets of ETS1 and ETS2. Of note, we focused further analysis only on the regions occupied by both ETS1 and ETS2 because deletion of only *Ets1* or *Ets2* failed to elicit any observable phenotypic defect.

In the current study, although we examine cell cycle as a possible mechanism of aberrant angiogenesis due to the simultaneous loss of *Ets1* and *Ets2*, the exact events that trigger the M-phase arrest are yet to be determined. A number of genes were identified to also be downregulated in the DKO ECs but were not associated with any ETS1 or ETS2 peaks (data not shown). One possible explanation might be that these genes are indirect targets of ETS1 and ETS2 and their expression is governed by noncoding RNAs (an enriched subcategory in the ChIP-seq microarray overlap) which themselves are regulated by ETS1 and ETS2.

In summary, the genetic and biochemical evidence presented herein suggests that ETS1 and ETS2 regulate the cellular machinery within ECs responsible for maintaining cell cycle progression and cell survival required for angiogenesis.

## Experimental procedures

### Mice

The *Ets2*^*fl/fl*^ (8) mice were described previously. A tamoxifen inducible *Tie2-Cre-ER*^*Tam*^ (10) was used to induce Cre expression only in the endothelial cells of postnatal mice. *Ets1*^*fl/fl*^; *Ets2*^*fl/fl*^ mice were bred >10 generations into the FVB/N background. All other animals were maintained on a mixed 129/Sv × FVB/N background. All mice were genotyped by PCR. Primers and conditions used are available upon request. Use and care of mice in this study was approved by The Ohio State University Institutional Animal Care and Use Committee.

### Construction of targeting vectors

The two loxP sites were introduced encompassing the 4.5kb DNA fragment between the BamHI site in intron 6 and the BamHI site located after exon 8 at the 3′ end of the gene (see Figure S1A). A neomycin resistance cassette flanked by FRT sites was inserted before the 3′ BamHI site. The linearized targeting construct was introduced into embryonic stem cells by electroporation, and after neomycin selection, homologous recombinants were identified by PCR and Southern blot analysis. The neomycin cassette was removed by expression of Flp. *Ets1*^*fl/+*^ ES cells were injected into B6 blastocysts, and the resulting chimeras were used to establish the *Ets1*^*fl*^ strain.

### Matrigel plug angiogenesis assay

For subcutaneous Matrigel plug injections, 10-week-old control and inducible EC-DKO mice were administered 2mg of tamoxifen via IP for 5 consecutive days and put on a tamoxifen diet for the course of the experiment. Five days after the last injection, 350μl of ice-cold Matrigel containing 5 × 10^5^ MVT1 cells was injected slowly just below the skin in the flank region of the mice using a 28½ G needle. Plugs were harvested 5 days post-injection for subsequent histological studies.

### Isolation of endothelial cells

Mouse aortic ECs were isolated as previously described (31). Briefly, thoracic aortas were dissected from anesthetized mice and cut longitudinally into 1-to 2mm^2^ pieces. Explants (4 to 6) were placed in fibronectin (50μg/ml)-coated dishes. A small volume of complete EC culture media (Dulbecco’s modified Eagle medium (DMEM)–F12 supplemented with 20% heat-inactivated fetal bovine serum (FBS), penicillin-streptomycin, 30μg/ml endothelial cell growth supplement (Upstate Biotechnology), and 10U/ml heparin (Sigma-Aldrich)) was added to the dishes and placed in 37°C incubator at 5% CO_2_. Migrating ECs were observed within 2 to 3 days and on day 7 the cells were trypsinized and the EC population was maintained in complete EC culture media in 37°C incubator at 5% CO_2_.

### Microarray gene expression profiling and analysis

Endothelial cells used in this study were isolated from three independent sets of wild-type, *Ets1*^*-/-*^;*Ets2*^*fl/fl*^ and *Ets2*^*fl/fl*^ mice and infected with lentiviral GFP with or without Cre. RNA harvested from control and DKO ECs was utilized for gene expression profiling using the Affymetrix GeneChip Mouse Genome 430 2.0 array. Background correction and quantile normalization was performed to adjust technical bias, and gene expression levels were summarized by RMA method (32). A filtering method based on percentage of arrays above noise cutoff was applied to filter out low expression genes. A linear model was employed to detect differentially expressed genes. In order to improve the estimates of variability and statistical tests for differential expression, a variance smoothing method with fully moderated t-statistic was employed for this study (33). The expression data was analyzed by Gene Set Enrichment Analysis (GSEA) as described previously (11) with gene sets obtained from indicated categories in ToppGene suite (34).

### Lentiviral transduction

Infections were performed as described previously (30). Briefly, ECs (5 × 10^5^) were cultured overnight and infected with ecotropic lentivirus (pHAGE-IRES-GFP) vectors with or without PGK-Cre. The infected cells were passaged or utilized for subsequent experiments 72 hours post-infection.

### Quantitative real-time PCR

Total RNA was extracted from cultured ECs using TRIzol (Invitrogen) according to the manufacturer’s instructions. Taqman probes (Applied Biosystems) were utilized to perform the qPCR. Normalized expression of genes compared to the reference ribosomal L4 gene was computed using a variation of the ddCt method (35).

### Histology and immunostaining

For Matrigel plugs, 5μm sections were prepared from harvested frozen plugs to perform immunostaining. Cultured aortic ECs plated on chamber slides were used for phospho-histone-H3 staining. Primary antibodies used were rat α-mouse CD31 (1:50 dilution; BD Biosciences) and rabbit α-mouse phospho-Histone H3 (Ser10) (1:100 dilution; Millipore). Alexa fluor 488 and 594 conjugated secondary donkey α-rat or donkey α-rabbit antibodies (1:250 dilution; Invitrogen) were used for fluorescent detection. A Nikon Eclipse E800 epifluorescence microscope equipped with a Photometrics Coolsnap camera and Nikon Plan Fluor 4x (N.A. 0.13), 10x (N.A. 0.30), 20x (N.A. 0.50), 40x (N.A. 0.75) and 100x (oil N.A. 1.3) objectives were used to acquire immunofluorescent images. MetaVue software from Molecular Devices was used for image acquisition. An Olympus FV1000 Filter Confocal system using a UPLFLN 40x oil objective (N.A. 1.3) was utilized for confocal microscopy. All images were acquired at room temperature. Blood vessel size and branching (Matrigel plug sections) were measured using the ‘connected region’ plugin and the ‘Analyze skeleton’ plugin, respectively, in FIJI.

### DNA content analysis and detection of apoptosis

Apoptosis was induced in cultured ECs through serum starvation (0.1% serum, 24 hours). After the induction, both adherent and non-adherent cells were harvested, pelleted, washed, and pelleted again. Cells were then subjected to PI (Roche) staining or Annexin V-Pacific Blue/7-AAD (Invitrogen/eBioscience) staining followed by fluorescent activated cell sorting (FACS) to assay apoptosis. Labeled cells were analyzed using a BD LSR II Flow Cytometer. Cells demonstrating DNA content less than that of G_0_/G_1_-phase cells were considered to be apoptotic (sub-G_0_). For Annexin V-Pacific Blue/7-AAD staining, cells positive for annexin V were considered to be apoptotic.

### Tube formation assay

A Matrigel tube formation assay was performed to characterize the vessel-forming ability of cultured ECs *in vitro.* 2 × 10^4^ cells were plated on a single layer of Matrigel and cultured in compete EC media. Development of vessels was monitored every 4 hours for 24 hours.

### Chromatin immunoprecipitation assays

Chromatin immunoprecipitation (ChIP) assays were performed on cultured aortic ECs as described previously (36). The chromatin was immunoprecipitated with 5μg anti-ETS1, 20μg anti-ETS2 antibodies, as characterized previously (8), and 20μl anti-Acetyl-CBP (Lys1535)/p300 (Lys1499) antibody (Cell Signaling). Immunoprecipitates were analyzed by real-time PCR using the Roche Universal Probe Library (Roche Diagnostics) and the Universal Master Mix (Applied Biosystems). Chip-sequencing was performed to combine chromatin immunoprecipitation (ChIP) with high throughput DNA sequencing to identify the binding sites of nuclear proteins in a genome-wide manner. Wild-type cultured aortic ECs were used to generate the immunoprecipitated and input control libraries using TruSeq DNA Sample Prep Kit v2 (Illumina) following manufacturer’s direction. The libraries were sequenced on a HiSeq 2000 system (Illumina).

### ChIP-seq analysis

Bowtie2 software (13) was used to map the sequence reads to the mm10 mouse genome. Peak calling and motif analysis were done with HOMER v3 software (14). Global annotation of peaks with respect to TSS and gene body was performed with the R Bioconductor package ChIPseeker (37). Centered K-means clustering was performed using Cluster 3.0 (21) and visualized using Java-Treeview program (23).

Over layered ChIP-seq peaks at noted loci were visualized using Genome browser in the Box (GBiB) (38). All external ChIP-seq datasets were either downloaded from GEO omnibus or European nucleotide archive as fastQ file format and processed as described above.

## Acknowledgements

The authors are grateful to the core facilities involved in this work, OSU ULAR and OSU-CCC Solid Tumor Biology Histology Core for tissue preparation. This study was supported by the NIH-NCI grant PO1CA097189 (MCO), NIH-NIAMS grant 2R01AR044719-15A (support for SSM), and NIH-NCI grant T32CA193201 (support for CBM).

